# Ventromedial Prefrontal Cortex and Basolateral Amygdala Projections to Ventromedial Striatum Encode Active Avoidance Behavior

**DOI:** 10.1101/2023.09.15.558002

**Authors:** Teagan E. Bullock, Lisa A. Gunaydin

**Affiliations:** Department of Chemistry San Francisco State University, San Francisco CA 94132; Department of Psychiatry and Behavioral Sciences University of California San Francisco, San Francisco CA 94158

## Abstract

Active avoidance is a fundamental defensive behavior, defined as performing a voluntary action to escape a noxious stimulus. Active avoidance serves as an adaptive response to threat, ensuring safety. In individuals with anxiety disorders and post-traumatic stress disorder, avoidance behavior can become maladaptive when even safe situations are avoided, at great psychosocial cost. Corticostriatal and amygdalostriatal pathways have been implicated in active avoidance, though their role in learning and expression of this behavior is not fully understood. Projections from the ventromedial prefrontal cortex (vmPFC) to the ventromedial striatum (VMS) have been shown to regulate a variety of defensive behaviors; however, their role in active avoidance has not previously been examined. Additionally, basolateral amygdala (BLA) projections to the VMS have been shown to promote active avoidance in studies utilizing pharmacological inactivation and optogenetic inhibition, though in vivo real time neural activity in this pathway has not been recorded during active avoidance learning. Here we utilized fiber photometry for in vivo recordings of neural activity in vmPFC-VMS and BLA-VMS projections during active avoidance learning and expression. We implemented a two-way signaled active avoidance paradigm in which a light cue served as the conditioned stimulus (CS) signaling an impending foot shock. We examined changes in neural activity in these two projections during CS presentation, onset of active avoidance, and conditioned freezing. We found that vmPFC-VMS projections develop learning-related increases in activity at CS onset across training, while BLA-VMS projections do not show learning-related encoding of the CS. Additionally, we found that both vmPFC-VMS and BLA-VMS projections develop an increase in activity at avoidance onset. No changes in neural activity were observed during cued freezing in either vmPFC-VMS or BLA-VMS projections. Together these results indicate that vmPFC-VMS projections encode both CS and avoidance, while BLA-VMS projections may simply encode avoidance. Finally, we utilized optogenetic inhibition of vmPFC-VMS projections during the CS to investigate the necessity of this pathway for expression of active avoidance behavior. We found that inhibition of vmPFC-VMS projections attenuates learned active avoidance behavior, indicating that activity in this pathway is required for proper active avoidance. In summary, our results demonstrate task-relevant encoding of active avoidance behavior in vmPFC-VMS and BLA-VMS projections.

## Introduction

Active avoidance is defined as performing a voluntary action to escape a noxious stimulus. This defensive behavior is highly conserved across species and is critical to survival in mediating adaptive response to threat. Avoidance behavior becomes maladaptive when it persists in inappropriate contexts, impairing normal functioning. Maladaptive avoidance is a key behavioral symptom in Post-Traumatic Stress Disorder (PTSD) and anxiety disorders. In the context of a perceived psychological threat, humans learn to engage in active avoidance in response to a “triggering” stimulus associated with anxiety or a traumatic experience. By performing a voluntary action, they can escape the perceived threat and the associated psychological distress. In maladaptive avoidance, repetitive engagement in this behavior leads to a cycle of reinforcement in which escaping the threat temporarily provides a feeling of safety, perpetuating future avoidance behavior (Borkovec et al., 1999). This can also lead to reinforcement of fear by preventing exposure to the trigger, eliminating any opportunity to unlearn the association between trigger and threat. (Lovibond et al., 2009; Salkovskis et al., 1999). Thus, where avoidance is manifested in neuropsychiatric disease, it is often a persistent symptom. In order to develop innovative treatments for these conditions, advancing our understanding of the neural circuitry encoding active avoidance learning and expression is critical.

Human functional Magnetic Resonance Imaging (fMRI) studies have implicated dysregulated ventromedial prefrontal cortex (vmPFC) amygdala, and ventral striatum (VS) activity in anxiety and PTSD. Disrupted functional connectivity between the amygdala and vmPFC is believed to contribute to behavioral symptoms of anxiety and PTSD (Kim et al., 2011; Myers-Schulz and Koenigs, 2014; Rauch et al., 2006). Additionally, functional synchronization in the neural circuit incorporating the mPFC, amygdala and striatum is necessary for adaptive responses to threat (Collins et al., 2014). In the few studies that examine these regions in avoidance specifically, activity in the vmPFC and amygdala is correlated with avoidance while activity in VS is correlated with aversive stimuli in both avoidance and unavoidable contexts (Jensen et al., 2003; Sun et al., 2020). However, human fMRI studies examining broad changes in neural activity in whole brain regions are not able to distinguish among specific neuronal subpopulations, limiting the interpretation of these results. Demonstrating functional heterogeneity among these subpopulations requires the use of high-resolution methods to dissociate specific neural projections forming key pathways in active avoidance circuitry. The limitations of clinical research in human subjects motivates circuit-level mechanistic studies in rodent models, which have further implicated vmPFC, BLA and VMS in active avoidance learning and expression.

Studies utilizing pharmacological inactivation of whole vmPFC, BLA, and VS yield mixed results. In some instances, pharmacological inactivation of whole BLA or whole VS was shown to impair avoidance while pharmacological inactivation of whole vmPFC did not impair avoidance (Bravo-Rivera et al., 2014). However, this opposed previous studies which suggested that pharmacological inactivation, delivery of protein synthesis inhibitor, and lesions to vmPFC all attenuated active avoidance (Moscarello and LeDoux, 2013). This discrepancy may be the result of methods that cannot distinguish among neural subpopulations within vmPFC. Thus, determining whether vmPFC activity is necessary for active avoidance requires more precise methods for delivering targeted inhibition to specific projection-defined vmPFC subpopulations.

Acquisition of active avoidance behavior is known to rely on a complex learning process incorporating both Pavlovian and instrumental learning (LeDoux et al., 2017). In a two-way active avoidance paradigm, mice are placed in a chamber and exposed to paired light and shock stimuli (Kyriazi et al., 2018). In Pavlovian learning, mice associate the conditioned stimulus (CS) light with the threat of an unconditioned stimulus (US) shock, and eventually the light alone is sufficient to induce a threat response (LeDoux et al., 2017). Following acquisition of the CS-US association, the primary threat response is passive freezing. In instrumental learning, the mouse learns to shuttle out of the shock chamber in response to the light cue in order to avoid shock. This behavior is reinforced with repetitive engagement, and active avoidance becomes the predominant threat response.

Selection of passive freezing or active avoidance (shuttling) behavior is determined by competition between their respective neural circuits (LeDoux et al., 2017). These opposing circuits are thought to be comprised of divergent output pathways from the basolateral amygdala (BLA) (Amaropanth et al., 2000; Ramirez et al., 2015; Jiminez & Maren, 2009). In rodent models, activity in the BLA to central amygdala (CeA) pathway promotes passive freezing (Jiminez & Maren, 2009; Janak & Tye, 2015; Namburi et al., 2015; Goosens & Maren 2003). Conversely, activity in the BLA-VMS pathway is believed to promote active avoidance behavior (Ramirez et al., 2015; Diehl et al., 2020). The circuit with the greater activity may ultimately determine the behavioral response to threat, i.e., whether the subject will engage in passive freezing or active avoidance in response to the light cue.

The dorsomedial (dmPFC) and ventromedial (vmPFC) regions of the medial prefrontal cortex (mPFC) have been implicated in regulating amygdala-dependent defensive behaviors including avoidance (Diehl et al., 2020; Moscarello & LeDoux, 2013). The vmPFC specifically has been implicated in exerting top-down control over relevant neural circuits in the active avoidance paradigm. As passive and active avoidance circuits compete for behavioral control, vmPFC is thought to inhibit CeA to suppress freezing (Moscarello & LeDoux, 2013). In this way, vmPFC is hypothesized to shift the balance of neural activity in favor of active avoidance indirectly, by inhibiting the opposing passive freezing circuit. However, it remains unclear whether the regulatory role of vmPFC also includes upregulation of active avoidance circuitry to directly promote avoidance behavior. Specifically, the role of vmPFC-VMS projections in encoding and regulating active avoidance has not been demonstrated. Given that BLA-CeA and vmPFC-CeA pathways converge to promote and regulate passive freezing, it is possible that BLA and vmPFC projections to VMS converge with similar functional organization to promote and regulate active avoidance.

The present study examined the role of vmPFC-VMS and BLA-VMS projections in encoding learning and expression of active avoidance. We used retrograde viral approaches to target specific neural projections from vmPFC and BLA to VMS with a genetically encoded calcium indicator (GCaMP), which enabled in vivo projection-specific fiber photometry recordings throughout training in a two-way active avoidance paradigm. We used similar retrograde viral approaches for projection-specific optogenetic inhibition of vmPFC-VMS neurons in the active avoidance training paradigm. We examined neural activity during CS presentation and at onset of active avoidance and conditioned freezing. Both vmPFC-VMS and BLA-VMS projections showed encoding of active avoidance behavior with neural activity increasing at avoidance movement initiation, but neither projection showed any activity modulation during cued freezing. We found that vmPFC-VMS projections, but not BLA-VMS projections, showed encoding of the CS response, with learning-related increases in neural activity at CS onset developing across learning. Finally, we found that inhibition of vmPFC-VMS activity at CS onset attenuates learned avoidance behavior. Together, these results suggest that distinct subpopulations of projection neurons within vmPFC and BLA encode distinct aspects of active avoidance.

## Materials and Methods

### Animals

Animal cohorts consisted of both male and female wild-type C57BL6/J mice purchased from Jackson Laboratories. Animals were raised in normal light conditions on a 12:12 hour light/dark cycle and were given access to food and water ad libitum. All experiments were conducted in accordance with protocol established by the Institutional Animal Care and Use Committee at the University of California, San Francisco.

### Stereotaxic Surgery, Viral Injection, and Fiberoptic Cannula Implantation

Surgeries were performed at 8-12 weeks of age. Mice were anesthetized with 5.0% isoflurane at an oxygen flow rate of 0.8-1.0 L/min and transferred to a stereotactic apparatus (Kopf Instruments, Tujunga, CA, USA) with a heating pad, where anesthesia was maintained at 1.5-2.0% isoflurane throughout surgery. Respiration and toe pinch were monitored, and oxygen and anesthesia levels were adjusted accordingly. Lidocane (0.5%) was administered topically on the scalp before incision. Viruses were injected at a flow rate of 100nL/min using a 10uL nanofil syringe (World Precision Instruments, Sarasota, FL, USA) with a 33-gague beveled needle. Following completion of viral injection, a 10-minute delay allowed fluid to settle into the tissue before retracting the needle. Mice were given 0.2 mL each of ketoprophen (0.2 mg/mL) and slow-release buprenorphine (0.006mg/mL) via subcutaneous injection at time of surgery, and at 12 hours post-surgery. Mice were then transferred to a clean recovery cage atop a heating pad until fully recovered from anesthesia.

For fiber photometry recording from vmPFC-VMS and BLA-VMS projections, we unilaterally injected 1500nL and 500nL of AAV1-Syn-Flex-GCaMP6m (Addgene) into the vmPFC and BLA respectively, and 300nL of 1:1 CAV2-Cre (Institut de Génétique Moléculaire de Montpellier, Montpellier, France) /AAV1.hSyn-mCherry (UNC Vector Core) into the VMS. Injection coordinates (in millimeters relative to bregma) were as follows: vmPFC (+ 1.8 A/P, +-0.35 M/L, −2.8 D/V), BLA (−1.4 A/P, +-3.3 M/L, −4.9 D/V), VMS (+ 1.5 A/P, +-0.55 M/L, −4.7 D/V). Finally, a 400um-core fiber optic cannula metal ferrule was implanted in the vmPFC (2.5mm length) and BLA (5mm length). Implant coordinates were as follows: vmPFC (+ 1.8 A/P, +-0.35 M/L, −2.6 D/V), BLA (−1.4 A/P, +-3.3 M/L, −4.7 D/V). Mice in the photometry cohort were allowed 6 weeks for viral expression and recovery before beginning the active avoidance training paradigm.

Prior to photometry surgeries, test surgeries were performed to confirm that the signal being recorded from vmPFC implant was not contaminated with signal from BLA neurons, and vice versa. This would be an issue only in the case that BLA-VMS projection neurons labeled via retrograde tracing have axon collaterals projecting from BLA-vmPFC, or if the converse were true. Using the coordinates above, we unilaterally injected 1500nL or 500nL of AAV1-Syn-Flex-GCaMP6m (Addgene) into *either* the vmPFC or BLA respectively, and 300nL of 1:1 CAV2-Cre (Institut de Génétique Moléculaire de Montpellier, Montpellier, France) /AAV1.hSyn-mCherry (UNC Vector Core) into the VMS. Test subjects were allowed 6 weeks for viral expression before performing perfusions, histology, and microscopy to carefully examine GCaMP expression in vmPFC and BLA. It was determined that using the described retrograde tracing methods to label vmPFC-VMS and BLA-VMS projections did not result in significant labeling of axon collaterals terminating in the other brain region. Thus, the two signals recorded simultaneously from individual photometry implants in either vmPFC or BLA did not overlap.

For optogenetic inhibition of vmPFC-VMS projections, we bilaterally injected 1500nL of AAV5.EF1a.DIO.eNpHR3.0-eYFP.WPRE.hGH (Addgene) or AAV5-EF1a-DIO-eYFP (control virus) into the vmPFC and 300nL of 1:1CAV2-Cre/ AAV1.hSyn-mCherry into the VMS. Injection coordinates for IL and VMS were identical to the previous study. A 2.5mm, 200um Core, 0.39NA optical fiber with 1.25mm Ceramic Ferrule was implanted unilaterally in the vmPFC. Implant coordinates for the vmPFC were (+ 1.8 A/P, +-0.35 M/L, −2.4 D/V). Mice in the optogenetics cohort were allowed 8 weeks for viral expression and recovery before beginning the active avoidance training paradigm.

### Active Avoidance Behavior

Subjects were run in a two-way active avoidance paradigm adapted from Kyriazi et al., 2018. Training occurred in a custom-made apparatus consisting of two connected chambers each with infrared and visible spectrum LED lights underneath a shock floor. Trials were conducted in the dark with infrared lights used to track mouse location using Ethovision XT (Noldus, Wageningen, Netherlands) video recording software. LED light (CS) and shock (US) presentations were controlled by an Arduino using custom-written code (Arduino, Somerville, MA, USA) with tracking data being communicated to Arduino continuously.

Before each session, subjects were placed in the chamber and allowed time for chamber acclimation and collection of baseline data. The duration of this acclimation period was 5 minutes on day 1 of training and 1 minute on all subsequent days of training. After each session, subjects remained in the chamber for 1 minute following completion of the last trial for all days of training.

During training, subjects underwent 30 trials per day for 5 days. Each trial was followed by a variable inter-trial interval (ITI) with a duration between 20-30 seconds chosen randomly. For each trial, subjects were exposed to a 10 second light cue (CS) followed by 10 seconds of light paired with a 0.3mA shock (CS + US). Light and shock stimuli were presented in the chamber that the mouse was in at the start of each trial. Subjects were able to avoid exposure to the shock (US) by shuttling into the other chamber during the 10 second light only (CS) period and remaining there until cotermination of light and shock at the end of the 10s CS + US period. This was defined as a successful active avoidance trial.

### Fiber Photometry Recording

For fiber photometry recordings, in vivo calcium signal served as our proxy for neural activity in vmPFC-VMS and BLA-VMS projections. Projection-specific expression of GCaMP, a genetically encoded calcium indicator protein (GECI), resulted in calcium-dependent fluorescent emission. Photometry data were acquired using a custom-designed rig based on a previously described setup (Adhikari et al., 2015). The interface consisted of an RZ5P photometry processor and Synapse software (Tucker Davis Technologies, Alachua, FL, USA). This interface controlled a 4 channel LED driver (DC400, Thorlabs, Newton, NJ, USA), which controlled four fiber-coupled LEDs, two 470nm and two 405nm. The 470nm and 405nm LEDs were sinusoidally modulated at 210 Hz and 320 Hz, respectively. Light was delivered to vmPFC-VMS and BLA-VMS projections individually via two separate 4-port fluorescence mini-cubes (Doric Lenses). Each fluorescence mini-cube received input from one 470nm and one 405nm channel and delivered the combined 405/470nm LEF output through an individual 0.48nA 400uM fiber optic patch cable (Doric Lenses). For the vmPFC, a 2m patch cable was connected to a 1×1 rotary joint to facilitate movement and prevent tangling, which was in turn connected to another 1m patch cable delivering the LEF output. For the BLA, a single 2m patch cable delivered the LEF output directly. Patch cables were connected to their respective fiber optic cannulas via a ceramic mating sleeve (Thorlabs).

The 470nm LED was used to excite GCaMP, while the 405nm LED was used to control for artifactual fluorescence. The emitted light was simultaneously collected through the fiber optic patch cable and delivered back to the fluorescence mini cube, where it was output to a Photoreceiver, focused, and sampled at 60Hz. Light emission from the vmPFC was focused using a Visible Femtowatt Photoreceiver Module (model 2151, Newport, DC Low, 2×10^10 amplification) while light emission from the BLA was focused using a Fluorescence Detector Amplifier Module (Doric Lenses, DC, 2×10^10 amplification). Raw photoreceiver data were extracted and analyzed using custom scripts in Python. Raw data signals were then separated by demodulation based their respective LED channel’s modulation frequency. To synchronize data collection systems, a TTL pulse was delivered from Ethovision XT to the photometry interface at session start continuing in alternating 5 second on/off epochs for the duration of the session.

### Optogenetic Inhibition

For optogenetic inhibition of vmPFC-VMS projections, we suppressed neural activity using projection-specific expression of halorhodopsin (NpHR). Optogenetic inhibition was achieved using a 535nm laser (Shanghai Laser & Optics Century Co. LTD) to stimulate NpHR in vmPFC-VMS projection neurons. The laser was delivered through a 2m 0.39nA 200um core optical fiber patch cable (Doric Lenses) connected to the cannula via a ceramic mating sleeve (Thorlabs). Mice remained attached to the optical patch cable for the duration of the training session on days 1-5 to control for any effects introduced by the novelty of the cable, though optogenetic inhibition was only delivered on Day 5. Laser delivery was controlled by Master 8 (A.M.P.I) pulse generator, which was in turn controlled by an Arduino operating on a custom script. In the active avoidance paradigm, laser onset was aligned to light cue (CS) onset, delivering 10 seconds of continuous inhibition at 5mW laser power for the duration of the light (CS) only period.

### Perfusions and Histology

Following the conclusion of active avoidance training, mice were anesthetized with 5% isoflurane and received subcutaneous injection of a lethal dose (1.0mL) of a 10mg/mL ketamine/1mg/mL xylazine solution. Mice were then perfused transcardially using 10mL of 1X PBS followed by 10mL of 4% paraformaldehyde (PFA). Brains were extracted and fixed in 2mL PFA for 24 hours, then transferred to a 30% sucrose in 1X PBS solution for 48 hours. Brains were then frozen and sliced on a sliding microtome (Lecia Biosystems, Wetzlar, Germany) with individual slices being transferred to a well plate and preserved in cryoprotectant.

Slices were washed with Triton X-100 (Sigma-Aldrich) in a 1X PBS solution and mounted onto slides (Fisherbrand Superfrost Plus, ThermoFisher Scientific, Waltham, MA, USA). Slides were shielded from light to preserve flourescence and air-dried overnight. Once dried, 80uL of ProlongGold antifade reagent (Invitrogen, ThermoFischer Scientific) was pipetted onto each slide and a cover slip (Slip-rite, ThermoFischer Scientific) was applied. Covered slides were again shielded from light and air-dried over night before beginning microscopy. Viral injection, expression, and implant targeting were verified using fluorescence microscope (Leitz DMRB, Leica). Mice with exceptionally poor viral expression and/or inaccurate implant placement were excluded from the dataset, with the experimenter blind to the animals’ behavioral performance. Histology images were obtained using a Zeiss LSM-710 confocal microscope (Carl Zeiss Microscopy, Jena, Germany).

### Behavioral Data Analysis

Integrated data analysis was performed in a Pycharm CE (Jetbrains, Prague, Czechia) environment using custom-written scripts in Python adapted from Loewke et al., 2022. Movement was analyzed using Ethovision’s built-in movement detection software. The following movement detection settings were used to identify movement initiations: 10 sample averaging 2.25/2.0cm/s start/stop velocity threshold. Movement initiation and chamber crossing timestamps were extracted from raw data output by Ethovision. Chamber crossing timestamps were used to classify active avoidance trials as successful or unsuccessful. Movement initiation timestamps were used in identifying avoidance movement initiation (movements during the 10 second CS only period of successful trials) and calculating latency (time between CS onset and avoidance movement initiation. Freezing data were obtained from processing Ethovision video files using EZTrack open-source code (Pennington et al., 2019) with the following parameters: 9.0 motion cutoff, 1000 freezing threshold, and 25 sample (1 sec) minimum freeze duration. Freezing data were analyzed to extract freezing initiation timepoints and length of freezing bouts. Freezing onset timepoints were used to identify cued freezing in response to the CS (freezing during the 10 second CS only period of all trials). Duration of freezing bouts was used to quantify cued freezing behavior across days of training. Light (CS), shock (US) and ITI start/end timestamps were extracted from Arduino and correlated with movement and freezing data. Trials were classified as successful or unsuccessful based on the relative timing of CS onset, US onset, and chamber crossing. If the subject shuttled out of the chamber during the 10 second CS only period and did not re-enter the chamber during the CS + US period, exposure to shock was avoided and the trial was classified as successful active avoidance. If the subject did not shuttle out of the chamber during the 10 second CS only period and was exposed to shock or returned to the chamber during the CS + US period receiving late shock exposure, the trial was classified as unsuccessful. Subjects that did not reach 80% successful avoidance (24 out of 30 trials) on day 5 were excluded.

### Photometry Data Analysis

In analyzing photometry data, calcium signals for all channels were first aligned to the onset of the TTL pulse delivered at session start. The 405nm control signal was subtracted from the 470nm GCaMP signal to control for artifactual fluorescence. This corrected signal was then fit to a polynomial over time to normalize the data and correct for bleaching, yielding the normalized DF/F value. For each trial, the normalized DF/F signal was z-scored to a baseline period (last 10 seconds of the ITI) preceding CS onset. For ITI movements, the normalized DF/F signal was z-scored to a baseline period of 10 seconds preceding the movement initiation.

Z-scored DF/F signals were used in time-locking neural activity to CS onset (also split into successful/unsuccessful trials), avoidance movement initiation, ITI movement initiation, and initiation of cued freezing. For each event, a peri-event time histogram (PETH) was generated representing the average neural activity pattern across all occurrences of the event during the period of interest. PETH signals were quantified by calculating the average signal within the following time windows:

CS onset: Baseline (−1 to 0 sec), CS response (0 to 1sec)

CS onset successful vs unsuccessful: Baseline (−1 to 0 sec), CS response (0 to 1sec)

Avoidance movement initiation: Baseline (−10 to −8 sec), Movement (0 to 1 sec)

ITI movement initiation: Baseline (−10 to −8 sec), Movement (0 to 1 sec)

Freezing: Baseline (−2 to −1.5 sec), Freezing (0 to 0.5 sec)

Histograms showing distributions of movement velocity and duration were generated in PRISM using 1cm/sec and 1 second bin width, respectively.

Photometry data was excluded for subjects with unstable implants or mating sleeve disconnection that impaired recording of calcium signal.

### Optogenetics Data Analysis

For our optogenetics experiment, behavioral data were analyzed using a modified version of our integrated data analysis script. Movement, freezing, and training data were extracted and analyzed as outlined above. The experimental design required that no subjects be excluded based on performance, and thus the 80% successful avoidance exclusion criteria was not applicable.

### Statistical Analysis

Statistical analysis was performed using Prism 8 (Graphpad Software, San Diego, CA, USA). For photometry experiment data analysis, Repeated Measures One-Way ANOVA with Geisser-Greenhouse correction with Sidak and Tukey’s Multiple Comparisons Test, and Two-Way ANOVA with Sidak’s correction for multiple comparisons were used. For optogenetics experiment data analysis, two-way repeated measures ANOVA (assume sphericity, Sidak’s correction for multiple comparisons) was used.

### Data and Code Accessibility

All data and code from this study are freely accessible through contacting the corresponding author directly.

## Results

### Photometry subjects successfully acquire active avoidance behavior

To record endogenous neural activity in vmPFC-VMS and BLA-VMS projection neurons, we used a retrograde viral targeting approach to express GCaMP in a projection-specific manner (**Figure 1A, Extended Data Figure 1-1**). We then utilized fiber photometry to record calcium-induced fluorescence from these neurons, acting as a proxy for neural activity (**Figure 1B**). We recorded from both vmPFC-VMS and BLA-VMS projections simultaneously in all subjects throughout active avoidance training. Mice were trained in a two-way active avoidance paradigm in which a light cue (CS) signaled that a shock (US) was imminent (**Figure 1C**). Training consisted of one session per day for five days, with 30 consecutive trials per session.

**Figure 1.**
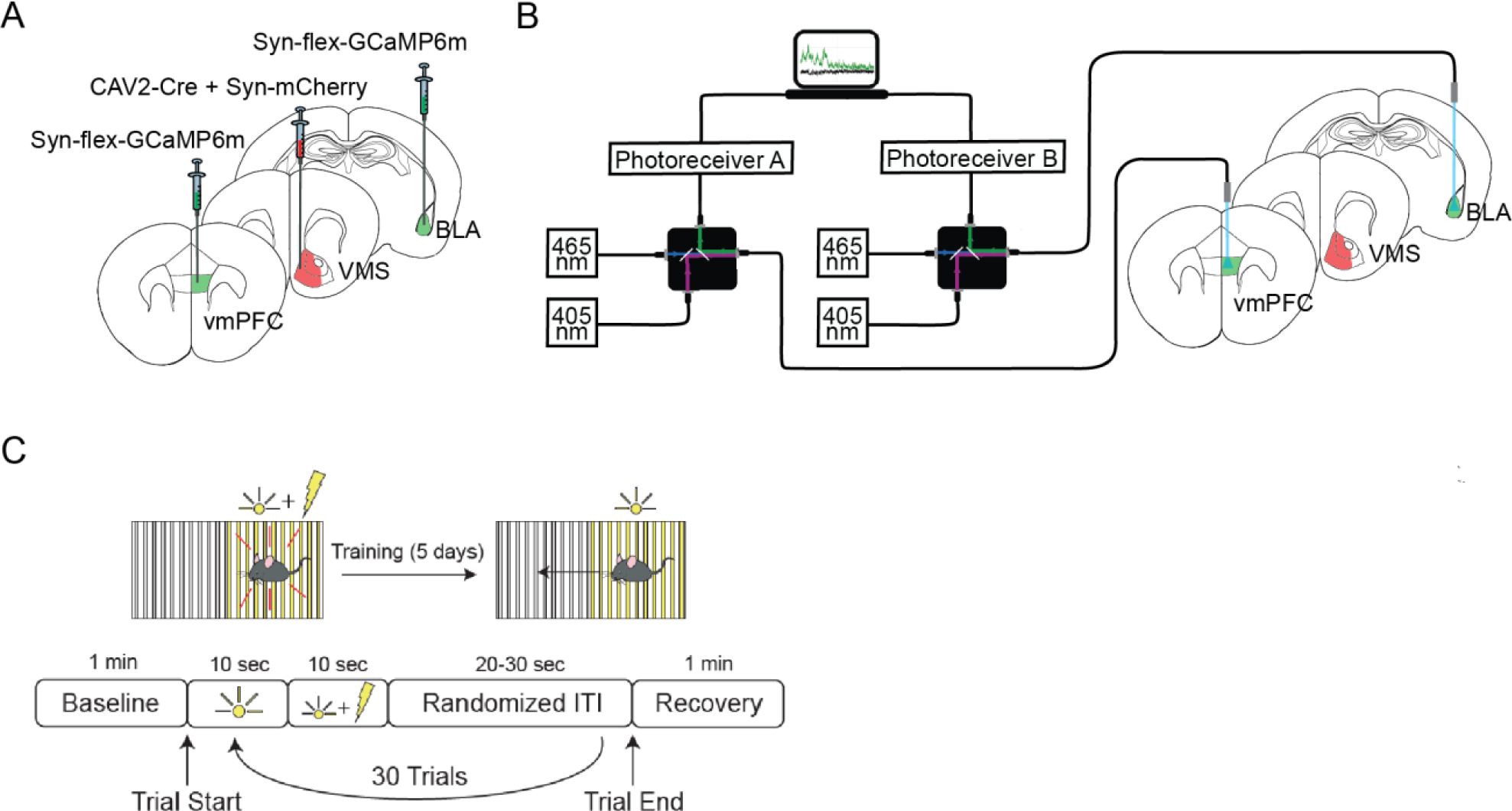
Fiber photometry recording from vmPFC-VMS and BLA-VMS projections during active avoidance training. (A) Viral targeting strategy for vmPFC-VMS and BLA-VMS fiber photometry. (B) Dual fiber photometry recording from vmPFC-VMS and BLA-VMS projection neurons expressing GCaMP6m. (C) Behavioral schematic for active avoidance training paradigm.

Throughout training, mice learned to shuttle out of the chamber in response to the CS to avoid exposure to shock. All subjects included in analysis reached 80% successful avoidance by Day 5 of training. Percent successful avoidance increased across training Days 1-5, while avoidance latency decreased (**Figure 2A**). Quantification of percent successful avoidance showed a significant increase in percent successful avoidance across Days 1, 3, and 5 of training (**Figure 2B**, One-Way Repeated Measures ANOVA with Geisser-Greenhouse correction, p < 0.0001 Tukey’s Multiple Comparisons Test, Day 1 vs Day 3 p < 0.0001, Day 1 vs Day 5 p < 0.0001, Day 3 vs Day 5 p = 0.0024, N = 9 mice). Quantification of avoidance latency showed a significant decrease in avoidance latency on Day 3 and Day 5 compared to Day 1, though no significant difference in latency was observed between Day 3 and Day 5 (**Figure 2C**, Repeated Measures One-Way ANOVA with Geisser-Greenhouse correction, Sidak’s Multiple Comparisons Test, Day 1 vs Day 3 p < 0.0001, Day 1 vs Day 5 p < 0.0001, Day 3 vs Day 5 p = 0.1638, N = 9 mice). These results indicate successful active avoidance learning and expression.

**Figure 2.**
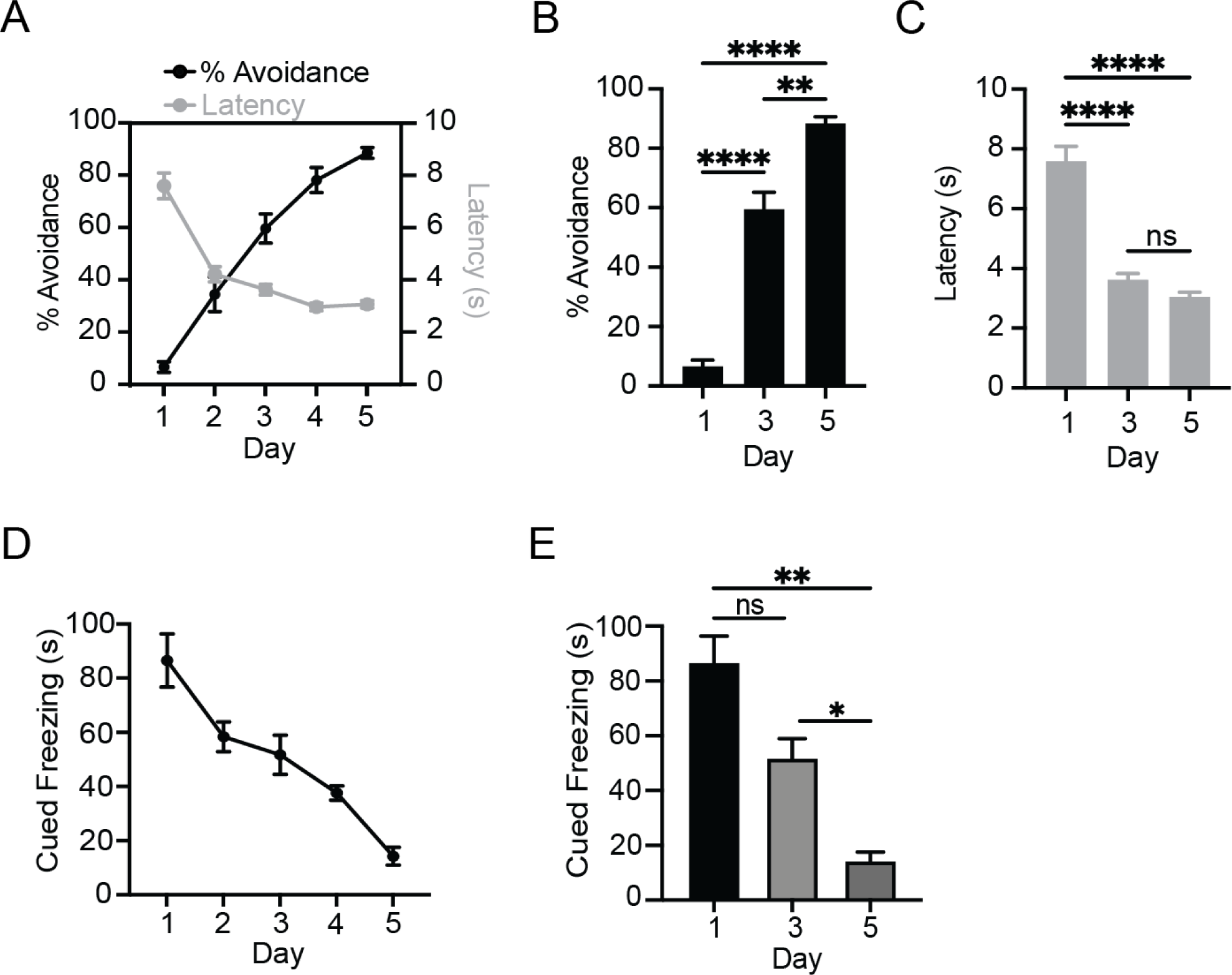
Avoidance increases across training while avoidance latency and cued freezing decrease. (A) Percent avoidance (number of successful trials out of 30) increases across training days while avoidance latency decreases. (B) Quantification of percent successful avoidance shows a significant increase in avoidance with learning. (C) Quantification of avoidance latency shows a significant decrease in latency with learning. (D) Cued freezing in response to CS decreases across training days with active avoidance learning. (E) Quantification of cued freezing shows cued freezing seconds are significantly lower on Day 5 compared to Day 1 and on Day 5 compared to Day 3. No significant difference in cued freezing seconds is shown on Day 3 compared to Day 1.

Cued freezing in response to CS decreased across training Days 1-5 (**Figure 2D**). Quantification of cued freezing seconds showed a significant decrease in cued freezing on Day 5 compared to Day 3 and Day 1, though no significant difference in cued freezing was observed between Day 1 and Day 3 (**Figure 2E**, One-Way Repeated Measures ANOVA, Geisser-Greenhouse correction, p = 0.003 Sidak’s Multiple Comparisons Test, Day 1 vs day 3 p = 0.4496, day 1 vs day 5 p = 0.0031, day 3 vs day 5 p = 0.0119). These results indicate suppression of passive freezing behavior with learning as active avoidance becomes the predominant behavioral response to CS.

### vmPFC-VMS and BLA-VMS projections show no change in activity during cued freezing

To examine changes in neural activity in vmPFC-VMS and BLA-VMS projections during cued freezing, we generated PETHs of z-scored change in calcium signal aligned to cued freezing onset (**Figure 3A**). Cued freezing was defined as freezing in response to CS during the 10 second CS-only period. This analysis used data from Day 1 as subjects demonstrated the greatest amount of cued freezing on this day. We found no significant change in calcium signal in vmPFC-VMS projections during the Freezing period in comparison to the Baseline period and no significant change in calcium signal in BLA-VMS projections during the Freezing period in comparison to the Baseline period (**Figure 3B**, Two-Way ANOVA, Task Period x Projection p = 0.6821, Task Period p = 0.9609, Projection p = 0.5325, Sidak’s Multiple Comparisons Test, vmPFC-VMS Freezing vs. vmPFC-VMS Baseline p = 0.9136, BLA-VMS Freezing vs. BLA-VMS Baseline p = 0.9415, N = 9 mice, n = 215 trials). These results suggest that vmPFC-VMS and BLA-VMS projections do not directly encode cued freezing behavior.

**Figure 3.**
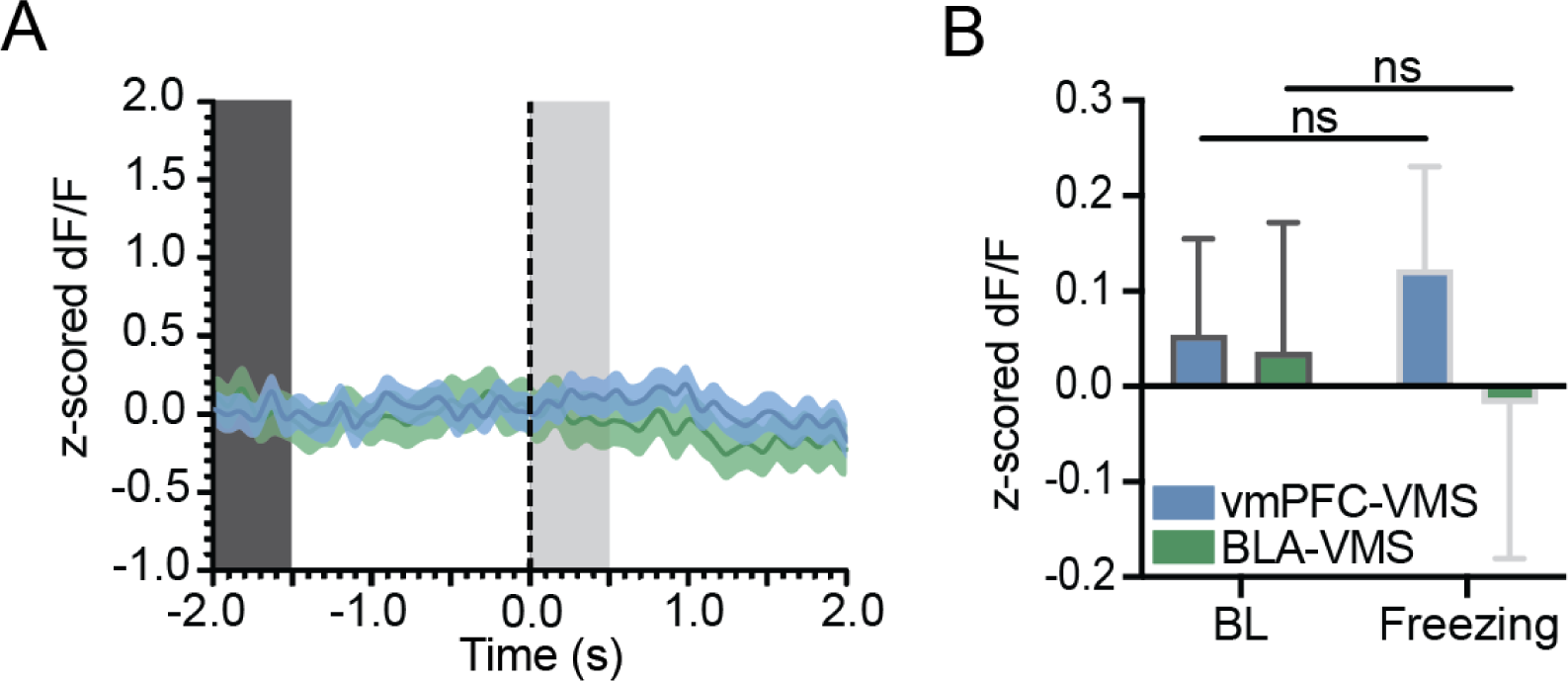
vmPFC-VMS and BLA-VMS projection neurons show no change in activity at onset of cued freezing. (A) Peri-event time histogram (PETH) showing no change in vmPFC-VMS calcium signal (z-scored DF/F) at initiation of freezing. Blue line: mean ± standard error of mean (SEM) for vmPFC-VMS; Green line: mean ± standard error of mean (SEM) for BLA-VMS; Dark grey box, baseline period (BL); light grey box, freezing period (Freezing). (B) Quantification of PETH aligned to initiation of freezing shows no significant change in calcium signal in vmPFC-VMS during freezing period (0 to 0.5s) compared to vmPFC-VMS during Baseline period (−2 to −1.5s), and no significant change in calcium signal in BLA-VMS during freezing period (0 to 0.5s) compared to BLA-VMS during Baseline period (−2 to −1.5s). ns = not significant, ** p **≤** 0.01.

### vmPFC-VMS and BLA-VMS projections show increased activity at avoidance onset

To examine changes in neural activity in vmPFC-VMS and BLA-VMS projections during avoidance behavior, we generated a PETH of z-scored change in calcium signal aligned to avoidance movement initiation (**Figure 4A**). This analysis used data from Day 5, in which the highest expression of active avoidance behavior was observed. We found that both vmPFC-VMS and BLA-VMS projections showed a significant increase in calcium signal during the Avoidance Movement period in comparison to the Baseline period (**Figure 4B**, Two-Way ANOVA, Task period x Projection p = 0.0225, Task period p < 0.0001, Projection p = 0.0162, Sidak’s Multiple Comparisons Test, vmPFC-VMS Baseline vs vmPFC-VMS Movement p < 0.0001, BLA-VMS Baseline vs BLA-VMS Movement p = 0.0452, vmPFC-VMS Baseline vs BLA-VMS Baseline p > 0.999, vmPFC-VMS Movement vs BLA-VMS Movement p = 0.0056, vmPFC-VMS N = 9 mice, n = 237 trials, BLA-VMS N = 7 mice, n = 182 trials). These results demonstrate encoding of active avoidance behavior initiation in both vmPFC-VMS and BLA-VMS projections.

**Figure 4.**
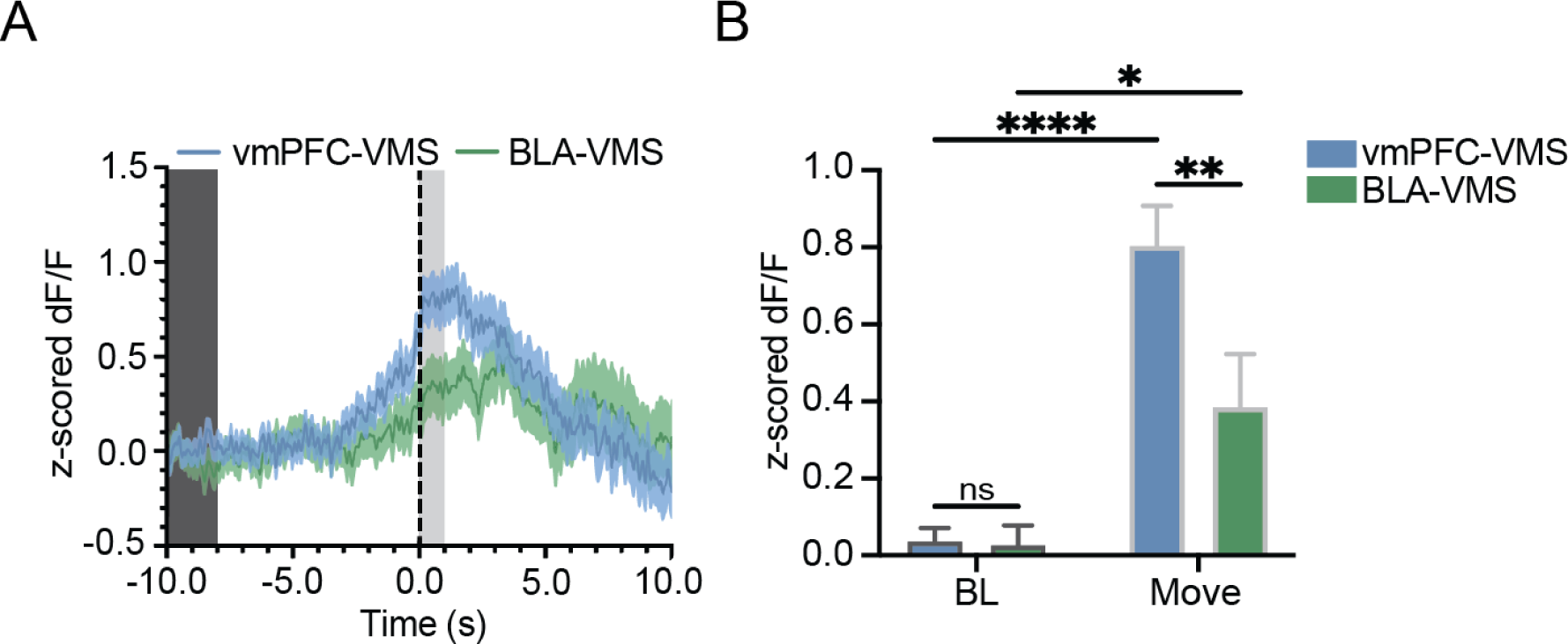
vmPFC-VMS and BLA-VMS projection neurons show increase in activity at avoidance movement initiation. (A) Peri-event time histogram (PETH) showing increase in vmPFC-VMS and BLA-VMS calcium signal (z-scored DF/F) at avoidance movement initiation. Blue line with shading represents mean ± standard error of mean (SEM) for vmPFC-VMS on Day 5. Green line with shading represents mean ± standard error of mean (SEM) for BLA-VMS on Day 5. Dark grey box, baseline period (BL); light grey box, avoidance movement period (Move). (B) Quantification of PETH aligned to avoidance movement initiation shows significant increase in vmPFC-VMS calcium signal during avoidance movement period (0 to 1s) compared to baseline period (−10 to −8s), significant increase in BLA-VMS calcium signal during avoidance movement period (0 to 1s) compared to baseline period (−10 to −8s, calcium signal in vmPFC-VMS projections is significantly higher than in BLA-VMS projections during the avoidance period (0 to 1s) and no significant difference in vmPFC-VMS and BLA-VMS projections during the baseline period (−10 to −8s). ns = not significant, * p **≤** 0.05, ** p **≤** 0.01, **** p **≤** 0.0001.

To determine whether these changes in neural activity were specific to avoidance movement initiation and not purely a result of general movement itself, we generated PETHs of z-scored change in calcium signal aligned to movement onset for non-avoidance movements occurring during the ITI (**Extended Data Figure 4-1**). We found that calcium signal for avoidance movements was significantly higher compared to ITI movements during the Movement Initiation period. To confirm that this difference was not a result of the unequal distributions of movement velocity and movement duration in avoidance and ITI movements, we compared movements of low velocity with movements of high velocity and compared movements of low duration with movements of high duration (**Extended Data Figure 4-2**). These results demonstrate that the increase neural activity in vmPFC-VMS and BLA-VMS projections during the Avoidance Movement Initiation period is not purely movement-related, and instead is specific to encoding active avoidance behavior.

### vmPFC-VMS projection shows learning-related increases in activity at CS onset

To examine learning-related changes in neural activity vmPFC-VMS projections in response to CS, we generated a PETH of z-scored change in calcium signal aligned to CS onset. This analysis used data from Day 1 and Day 5 to examine changes in neural activity before and after active avoidance learning. To isolate CS response, we examined the 1 second period of the CS only period immediately following CS onset, as the majority (> 85%) of avoidance movements occur after this time window (**Figure 5A**).

**Figure 5.**
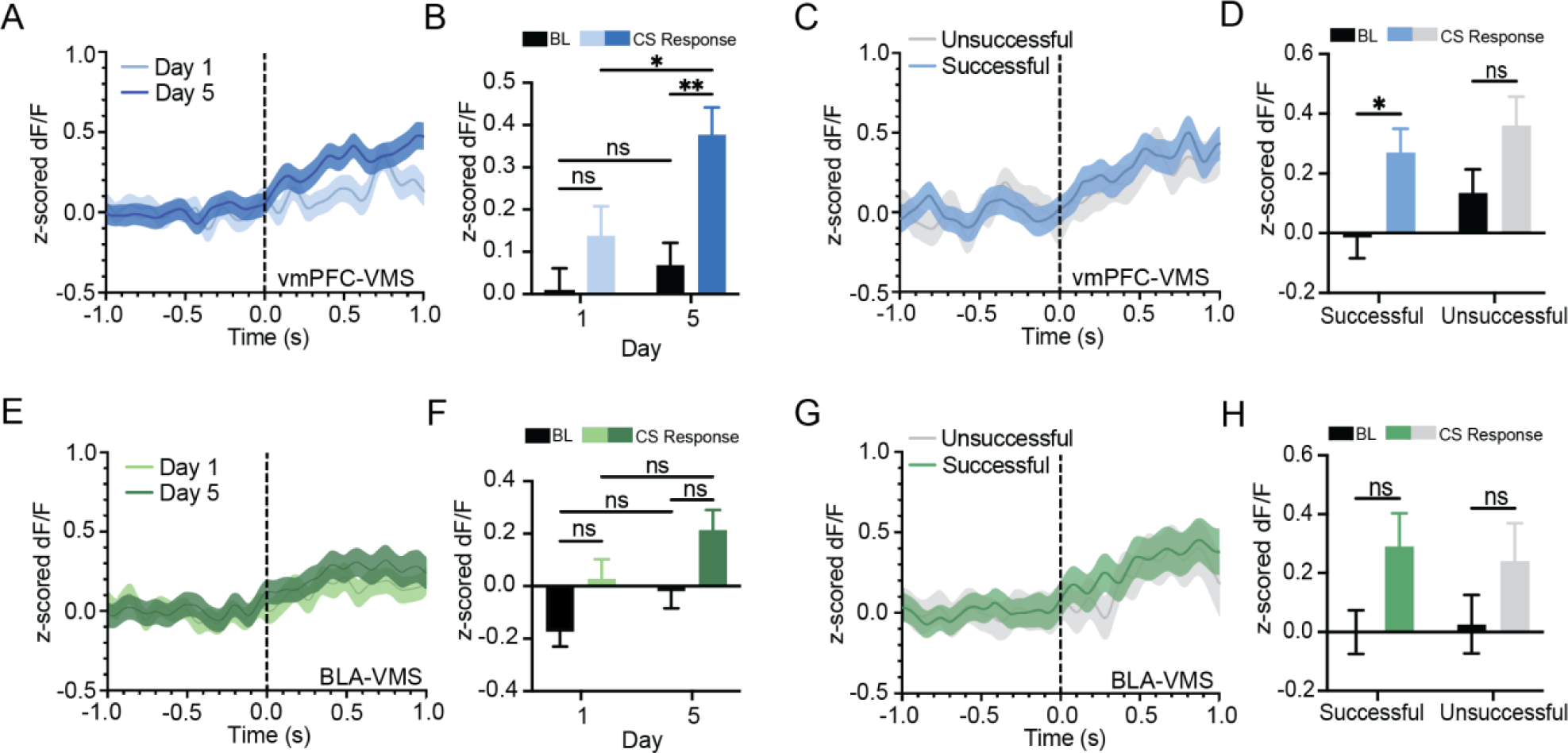
vmPFC-VMS, but not BLA-VMS, projection neurons show learning-related increase in activity at CS onset with training in active avoidance behavior paradigm. (A) Peri-event time histogram (PETH) showing increase in vmPFC-VMS calcium signal (z-scored DF/F) at CS onset. Light blue line, mean ± standard error of mean (SEM) for Day 1; dark blue line, mean ± standard error of mean (SEM) for Day 5. (B) Quantification of PETH aligned to CS onset shows vmPFC-VMS calcium signal is significantly higher during CS response period (0 to 1s) for Day 5 compared to CS response period for Day 1and compared to baseline period (−1 to 0s) for Day 5. No significant difference in calcium signal is observed between CS response period (0 to 1s) for Day 1 compared to baseline period (−1 to 0s) for Day 1. (C) PETH showing increase in vmPFC-VMS calcium signal at CS onset in successful trials (blue line) vs. unsuccessful trials (grey line). Data were obtained from Day 3 due to an equal number of successful and unsuccessful trials occurring on this day. (D) Quantification of PETH aligned to CS onset shows vmPFC-VMS calcium signal is significantly higher during CS response period (0 to 1s) compared to baseline period (−1 to 0s) in successful trials but not in unsuccessful trials. (E) PETH showing increase in BLA-VMS calcium signal at CS onset. Light green line, mean ± standard error of mean (SEM) for Day 1; dark green line, mean ± standard error of mean (SEM) for Day 5. (F) Quantification of PETH aligned to CS onset shows no significant difference in BLA-VMS calcium signal during CS response period (0 to 1s) compared to baseline period (−1 to 0s) for Day 5, for CS response period (0 to 1s) compared to baseline period (−1 to 0s) for Day 1, for CS response period (0 to 1s) for Day 5 compared to Day 1, or for baseline period (−1 to 0s) for Day 5 compared to Day 1.(G) PETH showing increase in BLA-VMS calcium signal at CS onset in successful trials (green line) vs. unsuccessful trials (grey line). (H) Quantification of PETH aligned to CS onset shows no significant change in BLA-VMS calcium signal during CS response period (0 to 1s) compared to baseline period (−1 to 0s) in successful trials or unsuccessful trials. ns = not significant, * p **≤** 0.05, ** p **≤** 0.01.

We found that vmPFC-VMS projections develop a learning-related peak in neural activity as CS onset, with a significant increase in calcium signal during the CS response period on Day 5 compared to the Baseline period on Day 5, and calcium signal during the CS response period on Day 5 being significantly higher compared to the CS response period on Day 1. No significant change in calcium signal was observed during the CS response period on Day 1 compared to the Baseline period on Day 1, and no significant change in calcium signal was observed in the Baseline period on Day 5 compared to the Baseline period on Day 1 (**Figure 5B**, Two-way ANOVA, Training Day x Task Period p = 0.1445, Training Day p = 0.0138, Task Period p = 0.0003, Sidak’s Multiple Comparisons Test, Day 1 Baseline vs Day 1 CS Response p = 0.5576, Day 5 Baseline vs Day 5 CS Response p = 0.0021, Day 1 Baseline vs Day 5 Baseline p = 0.9796, Day 1 CS Response vs Day 5 CS Response p = 0.0331, N = 9 mice, n = 270 trials). Taken together, these data suggest that encoding of CS response in vmPFC-VMS projections is learning-related, with an increase in neural activity in response to CS developing across active avoidance learning.

To further examine changes in neural activity in vmPFC-VMS projections in response to CS, we classified trials as successful and unsuccessful and split the data to generate a PETH of z-scored calcium signal aligned to CS onset for each trial type (**Figure 5C**). This analysis used data from Day 3 as there was a roughly equal number of successful and unsuccessful trials on this day. We found that vmPFC-VMS projections showed a significant increase in calcium signal during the CS response period for successful trials compared to the Baseline period for successful trials, and no significant change in calcium signal during the CS response period for unsuccessful trials compared to the Baseline period for unsuccessful trials (**Figure 5D**, Two-way ANOVA, Trial Type x Task Period p = 0.6832, Trial Type p = 0.2114, Task Period p = 0.0018, Sidak’s Multiple Comparisons Test, Successful Baseline vs Successful CS Response p = 0.0101, Unsuccessful Baseline vs Unsuccessful CS Response p = 0.1535, N = 9 mice, Successful n = 163 trials, Unsuccessful n = 107 trials). These data suggest that an increase in activity at CS onset in vmPFC-VMS projections is specifically associated with successful active avoidance.

### BLA-VMS projection shows no change in activity at CS onset

To examine learning-related changes in neural activity of BLA-VMS projections in response to CS, we generated a PETH of z-scored change in calcium signal aligned to CS onset (**Figure 5E**). We found that BLA-VMS projections show no significant change in calcium signal during the CS response period compared to the Baseline period for Day 1 or for Day 5. No significant change in calcium signal was observed in the CS response period on Day 5 compared to the CS response period on Day 1, and no significant change in calcium signal was observed in the Baseline period on Day 5 compared to the Baseline period on Day 1 (**Figure 5F**, Two-way ANOVA, Training Day x Task Period p = 0.8319, Training Day p = 0.0136, Task Period p = 0.0016, Sidak’s Multiple Comparisons Test, Day 1 Baseline vs Day 1 CS Response p = 0.1441, Day 5 Baseline vs Day 5 CS Response p = 0.1376, Day 1 Baseline vs Day 5 Baseline p = 0.5045, Day 1 CS Response vs Day 5 CS Response p = 0.3013, Day 1 N = 9 mice, Day 5 N = 7 mice, Day 1 n = 270 trials, Day 5 n = 210 trials).

To further examine changes in neural activity in BLA-VMS projections in response to CS, we classified trials as successful and unsuccessful and split the data to generate a PETH of z-scored calcium signal aligned to CS onset for each trial type (**Figure 5G**). We found that BLA-VMS projections show no significant change in calcium signal during the CS response period compared to the Baseline period for successful trials, and no significant change in calcium signal during the CS response period for unsuccessful trials compared to the Baseline period for unsuccessful trials (**Figure 5H**, Two-way ANOVA, Trial Type x Task Period p = 0.7169, Trial Type p = 0.9140, Task Period p = 0.0175, Sidak’s Multiple Comparisons Test, Successful Baseline vs Successful CS Response p = 0.0582, Unsuccessful Baseline vs Unsuccessful CS Response p = 0.3502, N = 9 mice Successful n = 163 trials, Unsuccessful n = 107 trials). Taken together, these data suggest that there is no encoding of CS response in BLA-VMS projections, with no learning-related changes in neural activity in response to CS developing across training, and no increase in neural activity in response to CS observed for either successful or unsuccessful trials.

### Optogenetic inhibition of vmPFC-VMS projections attenuates learned avoidance behavior

To inhibit activity in vmPFC-VMS projection neurons in vivo, we used a retrograde viral targeting approach to express NpHR in a projection-specific manner (**Figure 6A, Extended Data Figure 6-1**). We then delivered targeted inhibition to these neurons using a 535nm laser. Mice were trained in the same two-way active avoidance paradigm used for photometry recordings (**Figure 6B**). Optogenetic inhibition was delivered on Day 5 only, with laser onset aligned to CS onset and lasting for 10 seconds throughout the duration of the CS only period.

**Figure 6.**
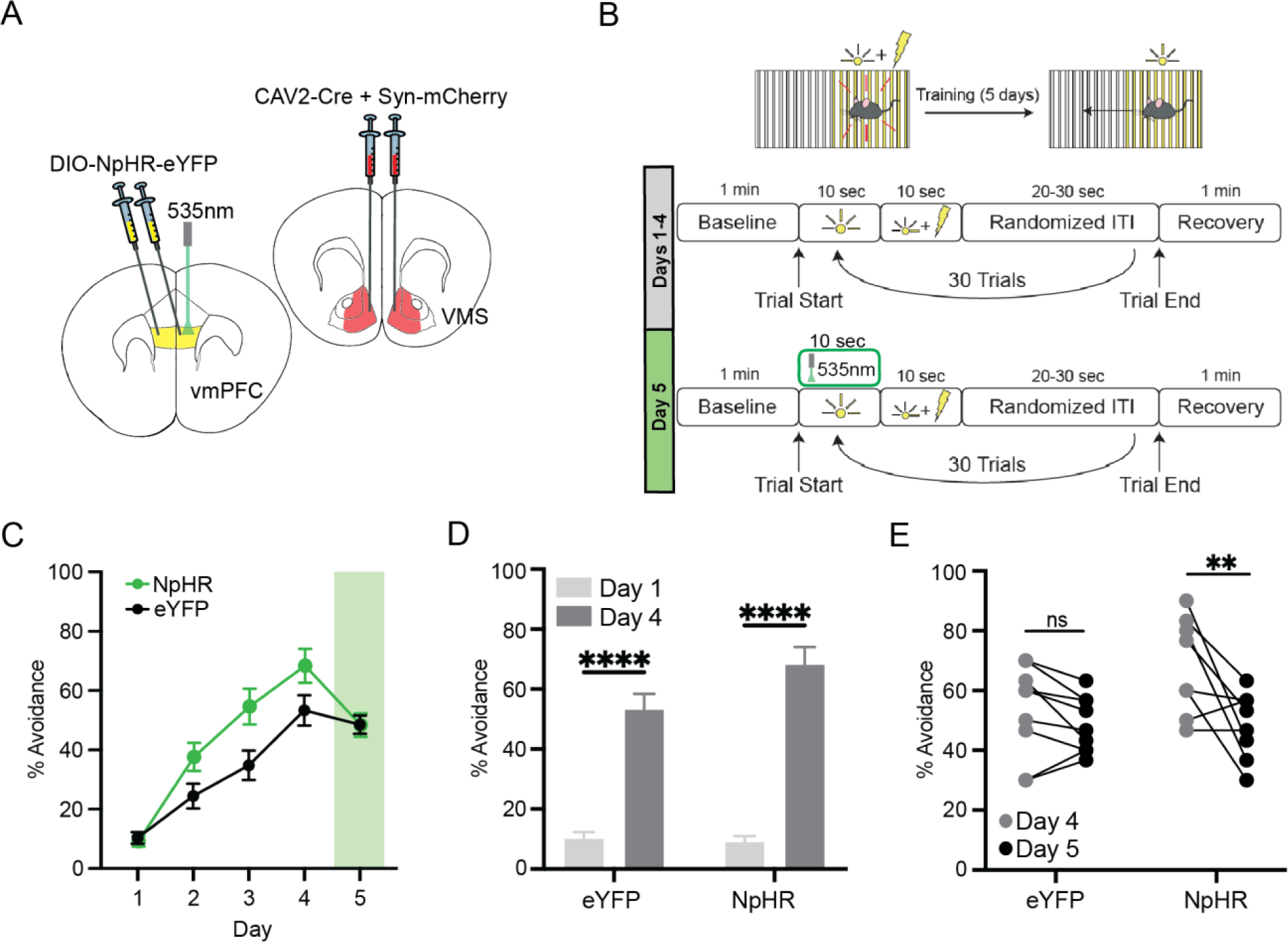
Optogenetic inhibition of vmPFC-VMS projection neurons attenuates learned active avoidance behavior. (A) Viral targeting strategy for vmPFC-VMS optogenetic inhibition. (B) Behavioral schematic for modified active avoidance training paradigm with optogenetic inhibition delivered at CS onset on day 5. (C) Percent avoidance (number of successful trials out of 30) increases with learning across training days 1-4 following the standard active avoidance training paradigm. Green bar indicates delivery of optogenetic inhibition on day 5. (D) Quantification of percent successful avoidance shows a significant increase on Day 4 compared to Day 1. (E) NpHR optogenetic inhibition group shows significant decrease in percent successful avoidance on Day 5 compared to Day 4. No significant difference in percent successful avoidance is observed on Day 5 compared to Day 4 for eYFP control group. ns = not significant, ** p **≤** 0.01, **** p **≤** 0.0001.

Percent successful avoidance increased across training Days 1-4 in both the NpHR group and eYFP control group (**Figure 6C**). Quantification of percent successful avoidance showed a significant increase in percent successful avoidance from Day 1 to Day 4 for both the NpHR group and eYFP control group (**Figure 6D**, Repeated measures two-way ANOVA, Condition x Training Day p = 0.1025, Condition p = 0.0572, Training Day p < 0.0001, Sidak’s Multiple Comparisons Test, eYFP Day 4 vs eYFP Day 1 p < 0.0001, NpHR Day 4 vs NpHR Day 1 p < 0.0001, eYFP N = 9 mice, NpHR N = 8 mice). These results indicate that learning occurred as normal across the first four days of training and active avoidance behavior was successfully acquired before delivery of optogenetic inhibition.

We found a significant decrease in percent successful avoidance with optogenetic inhibition on Day 5 compared to Day 4 in the NpHR group. No significant change in percent successful avoidance was observed on Day 5 compared to Day 4 in the eYFP control group. (**Figure 6E**, Repeated measures two-way ANOVA, Condition x Training Day p = 0.0563, Condition p = 0.1844, Training Day p = 0.0041, Sidak’s Multiple Comparisons Test, eYFP Day 5 vs eYFP Day 4 p = 0.5830, NpHR Day 5 vs NpHR Day 4 p = 0.0039, eYFP N = 9 mice, NpHR N = 8 mice). These results indicate that activity in vmPFC-VMS projections is necessary for expression of active avoidance behavior.

In summary, our results indicate that vmPFC-VMS and BLA-VMS projections play distinct roles in encoding active avoidance behavior. We found that neither vmPFC-VMS nor BLA-VMS projections encode cued freezing behavior, with no changes in neural activity observed at freezing onset. Additionally, we found that both vmPFC-VMS and BLA-VMS projections encode active avoidance movement initiation, with increases in neural activity at avoidance movement onset. Finally, we found that vmPFC-VMS projections, but not BLA-VMS projections, encode CS response, developing learning-related increases in activity at CS onset across active avoidance learning.

## Discussion

Our results implicate vmPFC-VMS projections in encoding the expression of active avoidance behavior during avoidance movement initiation. Furthermore, optogenetic inhibition of vmPFC-VMS projections attenuated learned avoidance behavior, suggesting that activity in this pathway is not only relevant but *necessary* for expression of active avoidance. Additionally, we found that vmPFC-VMS projections do not encode passive freezing behavior, suggesting that this neural subpopulation within the vmPFC preferentially encodes active as opposed to passive behavioral responses to threat. These neurons are therefore distinct from vmPFC projection neurons implicated in inhibiting CeA to suppress passive freezing (Moscarello & LeDoux, 2013). Thus, vmPFC-CeA and vmPFC-VMS pathways may serve complementary roles in suppression of passive freezing and promotion of avoidance behavior, respectively. In distinguishing these specialized neural subpopulations, we see that coordinated action selection in active avoidance depends on functional heterogeneity within the vmPFC. Given that the role of vmPFC in behavioral control is well established, it is likely that the primary role of vmPFC-VMS projections in active avoidance is to facilitate action selection when presented with a choice between the opposing behavioral responses of freezing and avoidance.

Furthermore, we found evidence of learning-related increases in activity in vmPFC-VMS projections in response to CS. As the CS signals the need for a choice between opposing behavioral responses, the role of vmPFC-VMS projections in action selection may be mediated by encoding of CS response in this pathway. Given that vmPFC-VMS activity is not necessary for expression of conditioned freezing which requires acquisition of association between CS and US, it is likely that learning-related increases in activity in vmPFC-VMS projections at CS onset are not explained by Pavlovian learning (Quirk et al., 2000). An alternate interpretation is that these learning-related changes are explained by vmPFC-VMS projections encoding another form of learning that is fundamental to active avoidance. That is, vmPFC-VMS projections may encode instrumental learning of the avoidance behavior in response to CS. This interpretation is consistent with research implicating vmPFC-VMS interaction in expression of CS-evoked instrumental behaviors (Keistler et al., 2015).

This interpretation is further supported by our finding that an increase in neural activity in vmPFC-VMS projections in response to CS distinguishes successful vs unsuccessful trials. Mid-way through the training paradigm on Day 3, the CS-US association is well established as evidenced by the presence of both avoidance and cued freezing in response to CS. The mix of successful (avoidance) and unsuccessful (freezing) trials suggests that on Day 3, the link between CS and avoidance behavioral response is still being acquired through instrumental learning, consistent with the two-factor theory of avoidance learning (LeDoux et al., 2017). If vmPFC-VMS projections were encoding Pavlovian learning, an increase in activity during the CS response period would be expected for all trials regardless of whether the subject engages in active avoidance (successful) or passive freezing (unsuccessful). However, increases in activity in response to CS were only observed when the subject successfully engaged in instrumental avoidance behavior. Thus, the development of learning-related increases in activity in vmPFC-VMS projections at CS onset may reflect instrumental learning and this encoding may mediate top-down regulation of instrumental actions to establish active avoidance as the predominant behavioral response.

In contrast, we found that neural activity in BLA-VMS projections encodes active avoidance behavior at avoidance movement initiation, though there is no evidence of encoding CS response or developing any learning-related changes in activity in response to CS. Considering the theory that CS information is processed in BLA before being transmitted to VMS to drive the avoidance response, our results suggest that BLA-VMS projection neurons represent a discrete neural subpopulation distinct from other subpopulations within BLA found to encode CS (Campeau & Davis, 1995; Maren, 1999; Fanselow & LeDoux, 1999; LeDoux et al., 2017). Additionally, these findings offer new perspective on the theory that the BLA-VMS pathway is most relevant to the instrumental learning component of active avoidance (Ramirez et al., 2015). Our results suggest that this pathway may not encode Pavlovian or instrumental learning components, but rather encodes execution of the avoidance behavior. Taken together, these results indicate that higher level cortical regions such as vmPFC-VMS projections may be responsible for integrating stimuli and selecting an appropriate behavioral response, while BLA-VMS projections may be more proximally involved in driving the behavior itself.

Additionally, we found that BLA-VMS projections do not encode passive freezing behavior. Considering studies implicating the BLA in expression of conditioned freezing (Jiminez & Maren, 2009; Janak & Tye, 2015; Namburi et al., 2015; Goosens & Maren 2003; LeDoux et al., 2017), these findings demonstrate functional heterogeneity of different BLA neuronal subpopulations in encoding different defensive behaviors (Amaropanth et al., 2000). Our results show that BLA-VMS projection neurons represent a neuronal subpopulation within the BLA encoding active avoidance behavior that is distinct from those subpopulations known to encode conditioned freezing behavior in response to threat. In contrast, activity in BLA-CeA projections is believed to drive expression of cued freezing (Jiminez & Maren, 2009; Janak & Tye, 2015; Namburi et al., 2015; Goosens & Maren 2003; LeDoux et al., 2017). Given that BLA-VMS and BLA-CeA projections are considered to be a key point of divergence in BLA in encoding of threat response, it is likely that the BLA-VMS projections that drive active avoidance directly oppose BLA-CeA projections encoding freezing (Amaropanth et al., 2000; Ramirez et al., 2015).

Incorporating the findings from this study in context with the larger body of research on the neural circuitry underlying active avoidance, we propose an integrated model (**Figure 7**). In the Pavlovian learning phase of active avoidance training, acquisition of CS-US association is encoded in distinct neuronal subpopulations within BLA (Campeau & Davis, 1995; Maren, 1999; Fanselow & LeDoux, 1999; LeDoux et al., 2017). In early training, activity in the BLA-CeA pathway is dominant, driving expression of conditioned freezing in response to CS (Jiminez & Maren, 2009; Janak & Tye, 2015; Namburi et al., 2015; Goosens & Maren 2003; LeDoux et al., 2017). In the instrumental learning phase, vmPFC-VMS projections may encode the instrumental learning of avoidance in response to CS. The balance of neural activity shifts from the BLA-CeA pathway to the BLA-VMS pathway, driving expression of active avoidance behavior.

**Figure 7.**
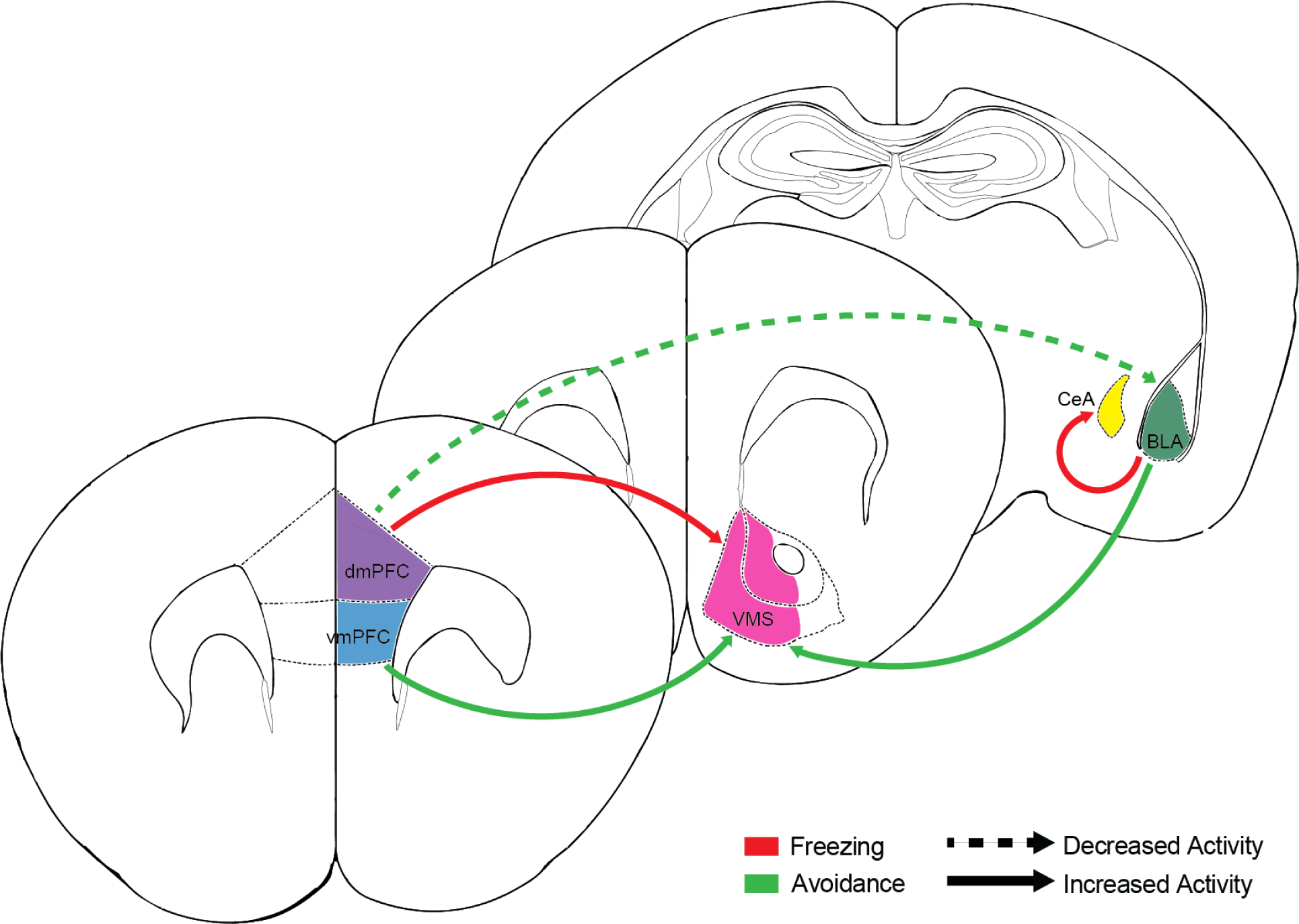
Proposed integrated model of relevant neural circuitry underlying active avoidance and passive freezing. Green arrows indicate projections that encode active avoidance. Red arrows indicate projections that promote encode passive freezing. Solid arrows indicate an increase in projection activity. Dashed arrows indicate a decrease in projection activity. Increased dmPFC-VS and BLA-CeA projection activity promotes freezing. Decreased dmPFC-BLA activity encodes active avoidance. Increased vmPFC-VMS and BLA-VMS projection activity encodes active avoidance but not freezing.

Both vmPFC and dmPFC exert top-down control over the relevant neural circuits throughout training, facilitating action selection to drive either passive freezing or active avoidance in response to CS. Another study showed that dmPFC-VS projections attenuate avoidance and promote freezing, while our results suggest that vmPFC-VMS projections promote avoidance over freezing (Diehl et al., 2020). Thus, converging corticostriatal pathways from dmPFC and vmPFC to VS may serve opposing roles in regulating different defensive behaviors. Furthermore, this opposition relies on similar functional organization of vmPFC and dmPFC projections to downstream brain regions within freezing and avoidance circuitry. Future studies should consider this functional organization in aiming to identify the role of other vmPFC projections in encoding active avoidance. Given that decreased activity in dmPFC-BLA projections encodes active avoidance (Kajs et al., 2022), vmPFC-BLA projections may also encode avoidance, potentially in an opposing manner. Furthermore, given that vmPFC has been found to inhibit dmPFC, projections between these two mPFC subregions may mediate their competition and thus reconcile action selection in choosing a given defensive behavior (Ji & Neugebauer, 2015).

Overall, our findings demonstrate that encoding of active avoidance in the vmPFC and BLA is more complex and heterogenous than previously thought. Within these regions, vmPFC-VMS and BLA-VMS projection neurons define distinct subpopulations specialized for active avoidance behavior, each with unique functions. We showed that the vmPFC-VMS pathway encodes both active avoidance movement initiation and CS response, while the BLA-VMS pathway encodes avoidance movement initiation but not CS response. Together, our results in the context of the broader avoidance literature suggest a role for amygdalostriatal projections in driving active avoidance behavior and opposing corticostriatal projections in regulating active avoidance versus passive freezing.

## Acknowledgments

We would like to acknowledge Dr. Adrienne Loewke and Dr. John Webb for providing original code used in integrated data analysis, and Dr. Pinelopi Kyriazi and Dr. Drew B. Headly from Dr. Dennis Pare’s lab for providing custom Arduino code for active avoidance training and for their guidance in constructing the active avoidance training apparatus.

---

**Extended Data Figure 1-1.**
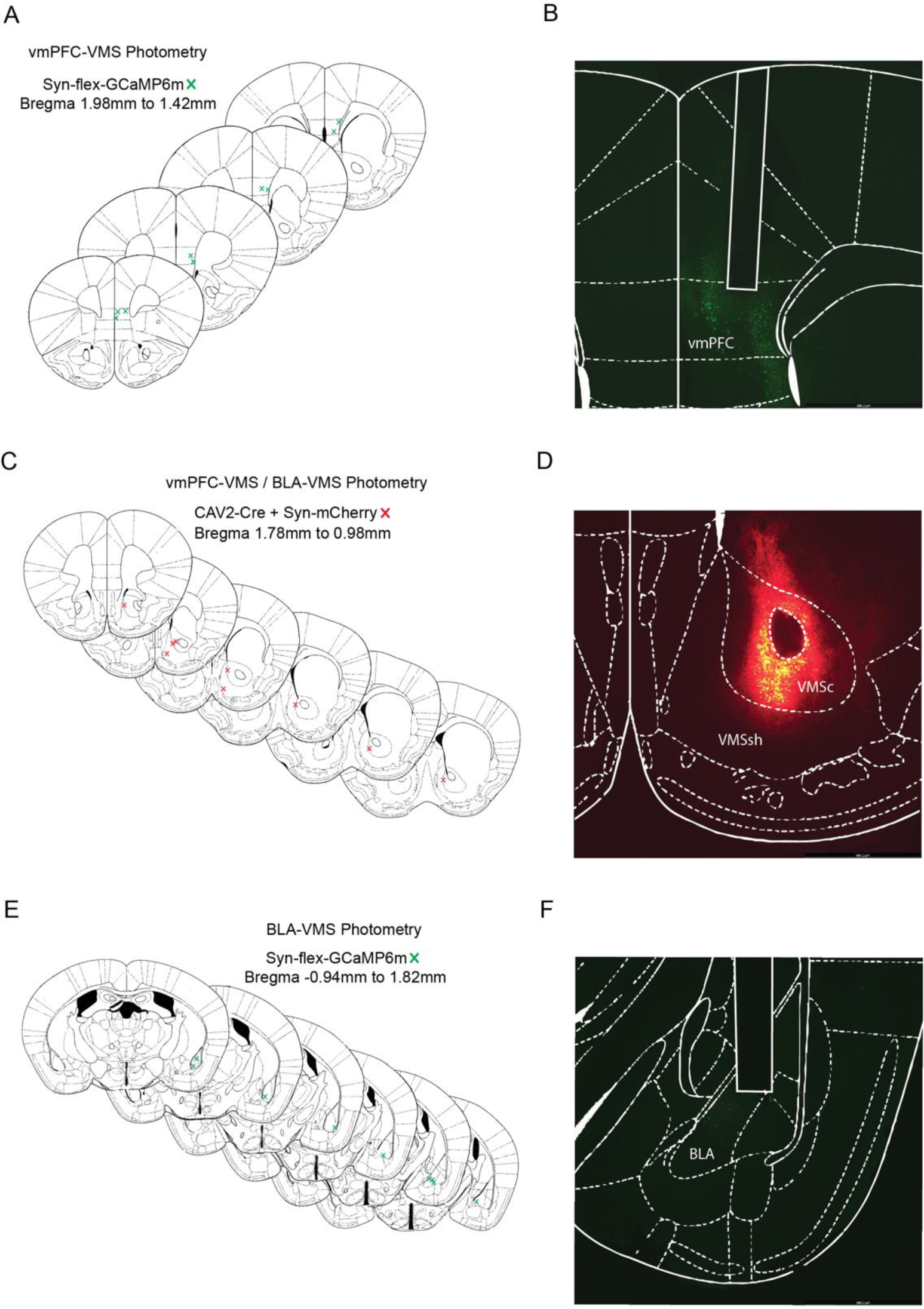
Histology and targeting for vmPFC-VMS and BLA-VMS photometry surgeries. (A) Verification of GCaMP viral injection in vmPFC (N = 9 mice). (B) Representative histological image of photometry implant and GCaMP viral expression in vmPFC. (C) Verification of CAV2-Cre + mCherry viral injection in VMS (N = 9 mice). (D) Representative histological image of CAV2-Cre + mCherry viral expression in VMS. (E) Verification of GCaMP viral injection in BLA (N = 9 mice). (F) Representative histological image of photometry implant and GCaMP viral expression in BLA.

**Extended Data Figure 4-1.**
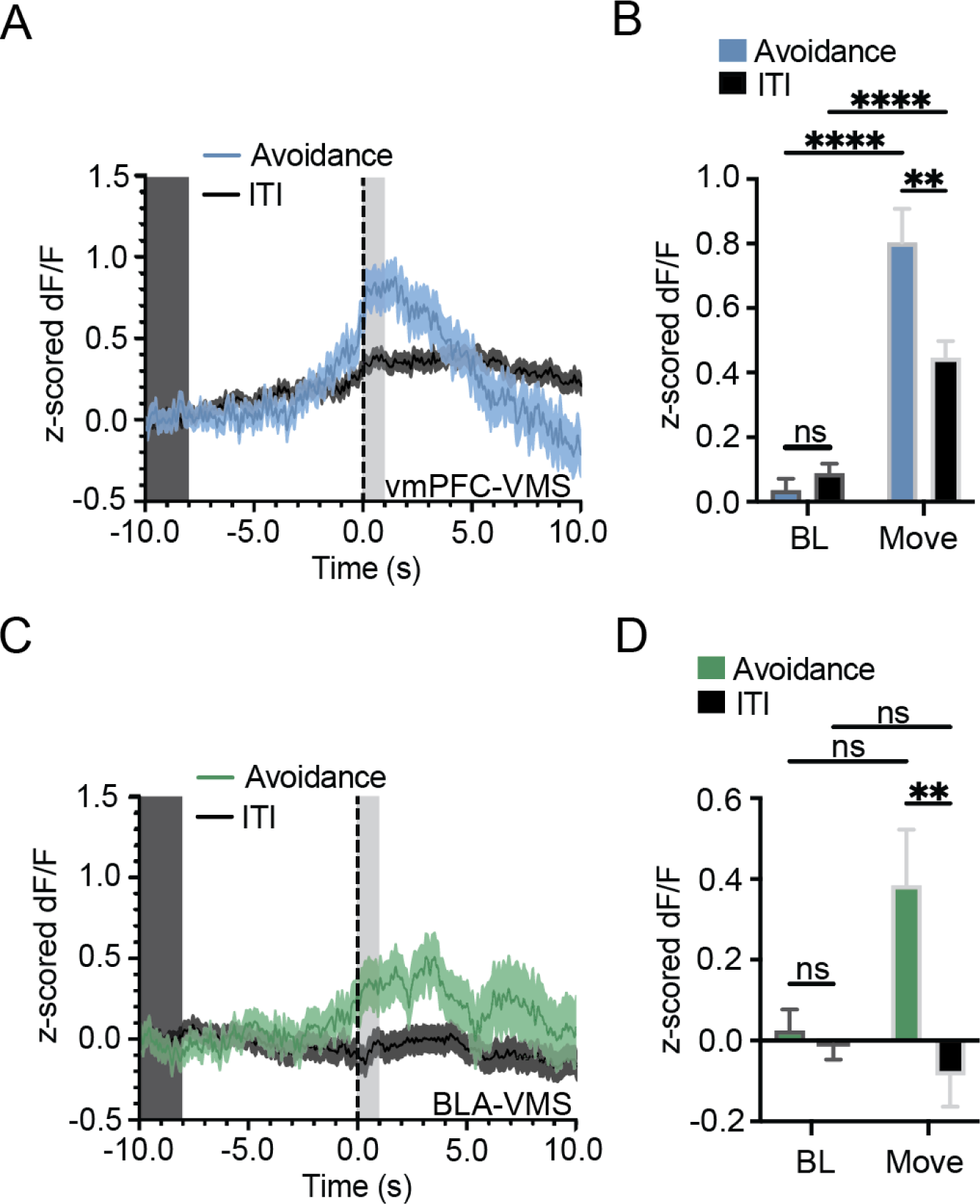
Increased activity in vmPFC-VMS and BLA-VMS projections during avoidance is not purely movement related. (A) Peri-event time histogram (PETH) showing increase in vmPFC-VMS calcium signal (z-scored DF/F) at movement initiation. Blue line, mean ± standard error of mean (SEM) for avoidance movements, black line, mean ± standard error of mean (SEM) for ITI movements. Dark grey box, baseline period (BL); light grey box, movement period (Move). (B) Quantification of PETH aligned to movement initiation shows vmPFC-VMS calcium signal is significantly higher during the movement period (0 to 1s) in avoidance movements compared to ITI movements, with a significant increase in calcium signal during movement period (0 to 1s) compared to baseline period (−10 to −8s) for both avoidance movements and ITI movements. (Two-way ANOVA, Task Period x Movement Type p = 0.0028, Task Period p < 0.0001, Movement Type p = 0.0265, Sidak’s Multiple Comparisons Test, Avoidance Baseline vs ITI Baseline p = 0.9949, Avoidance Baseline vs Avoidance Movement p < 0.0001, ITI Baseline vs ITI Movement p < 0.0001, Avoidance Movement vs ITI Movement p = 0.0014, N = 9 mice, Avoidance Movement n = 237 trials, ITI Movement n = 1184 occurrences). (C) PETH showing increase in BLA-VMS calcium signal (z-scored DF/F) at movement initiation. Green line, mean ± standard error of mean (SEM) for avoidance movements, black line, mean ± standard error of mean (SEM) for ITI movements. Dark grey box, baseline period (BL); light grey box, movement period (Move). (D) Quantification of PETH aligned to movement initiation shows BLA-VMS calcium signal is significantly higher during the movement period (0 to 1s) in avoidance movements compared to ITI movements, though there is no significant increase in calcium signal during movement period (0 to 1s) compared to baseline period (−10 to −8s) for either avoidance movements or ITI movements. (Two-way ANOVA, Task Period x Movement Type p = 0.0380, Task Period p = 0.1790, Movement Type p = 0.0156, Sidak’s Multiple Comparisons Test, Avoidance Baseline vs ITI Baseline p > 0.9999, Avoidance Baseline vs Avoidance Movement p = 0.3236, ITI Baseline vs ITI Movement p = 0.9258, Avoidance Movement vs ITI Movement p = 0.0089, N = 7 mice, Avoidance Movement n = 182 trials, ITI Movement n = 998 occurrences). ns = not significant, ** p **≤** 0.01, **** p **≤** 0.0001.

**Extended Data Figure 4-2.**
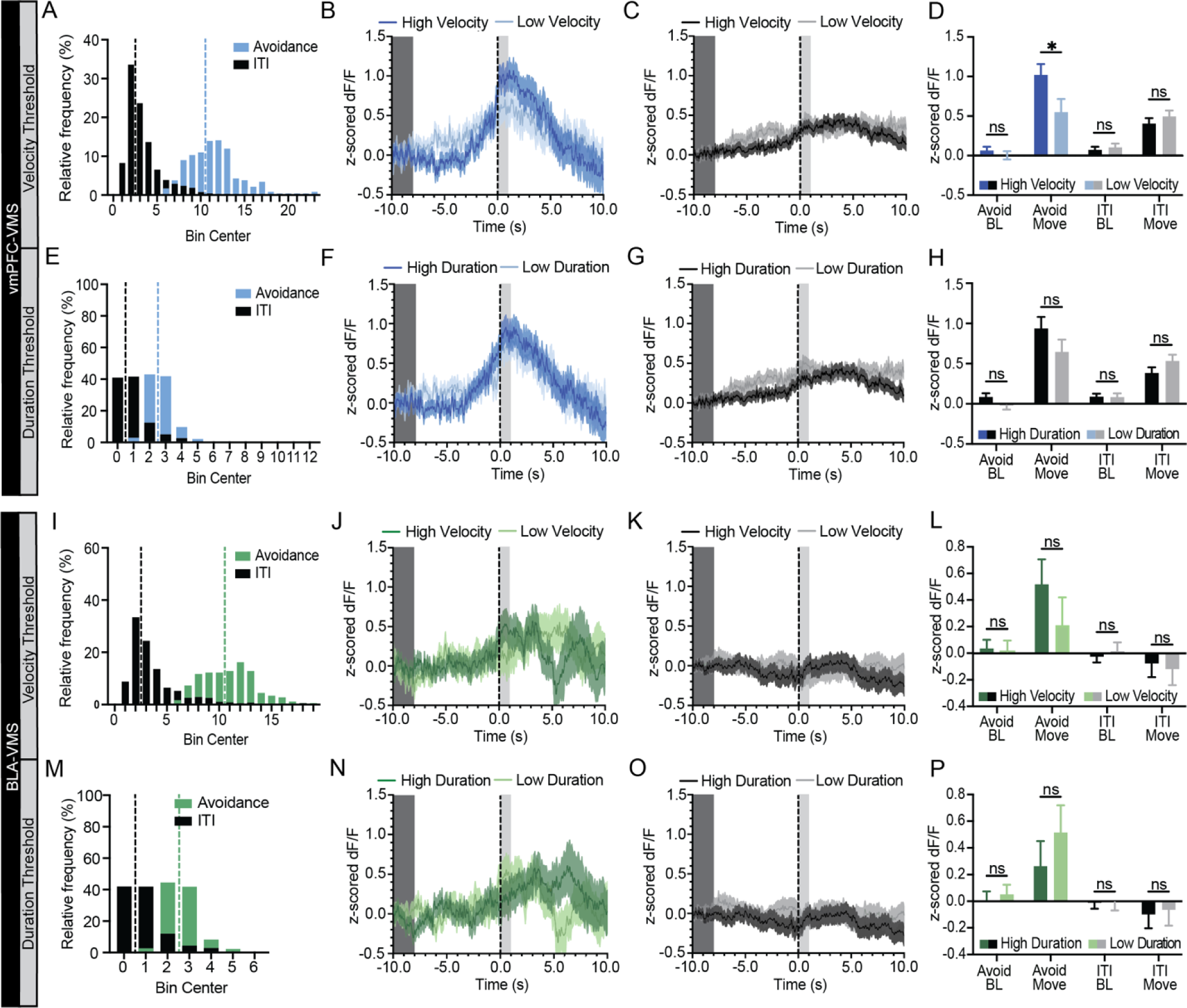
Differences in activity between avoidance and ITI movements is not a result of unequal distribution of movement velocity and duration. (A) Distribution of movement velocities for avoidance (blue) and ITI (black) movements for vmPFC-VMS data. Blue dashed line represents cutoff (10.5 cm/s) for separating avoidance movements of high and low velocity. Black dashed line represents cutoff (2.5cm/s) for separating ITI movements of high and low velocity. (B) Peri-event time histogram (PETH) showing increase in vmPFC-VMS calcium signal at avoidance movement initiation for high and low velocity avoidance movements. Light blue line, mean ± standard error of mean (SEM) for low velocity avoidance movements; dark blue line, mean ± standard error of mean (SEM) for high velocity avoidance movements Dark grey box, baseline period (BL); light grey box, movement period (Move). (C) PETH showing increase in vmPFC-VMS calcium signal at ITI movement initiation for high and low velocity ITI movements. Grey line, mean ± standard error of mean (SEM) for low velocity ITI movements; black line, mean ± standard error of mean (SEM) for high velocity ITI movements Dark grey box, baseline period (BL); light grey box, movement period (Move). (D) Quantification of PETH shows no significant difference in vmPFC-VMS calcium signal between high and low velocity movements for both avoidance and ITI movements during the baseline period (−10 to −8s) and for ITI movements during the movement period (0 to 1s). Avoidance movements show significantly higher vmPFC-VMS calcium signal during the movement period (0 to 1s) in high velocity compared to low velocity avoidance movements. (Two-way ANOVA, Task Period x Velocity p = 0.0373, Task Period p < 0.0001, Velocity p = 0.1357, Sidak’s Multiple Comparisons Test, High Velocity Avoidance Baseline vs Low Velocity Avoidance Baseline p = 0.9943, High Velocity Avoidance Movement vs Low Velocity Avoidance Movement p = 0.0333, High Velocity ITI Baseline vs Low Velocity ITI Baseline p = 0.9922, High Velocity ITI Movement vs Low Velocity ITI Movement p = 0.6998, N = 9 mice, High Velocity Avoidance Movement n = 134 trials, Low Velocity Avoidance Movement n = 100 trials, High velocity ITI Movement n = 683 occurrences, Low Velocity ITI Movement n = 493 occurrences). (E) Distribution of movement durations for avoidance (blue) and ITI (black) movements for vmPFC-VMS data. Blue dashed line represents cutoff (2.5 s) for separating avoidance movements of high and low duration. Black dashed line represents cutoff (0.5 s) for separating ITI movements of high and low duration. (F) PETH showing increase in vmPFC-VMS calcium signal at avoidance movement initiation for high and low duration avoidance movements. Light blue line, mean ± standard error of mean (SEM) for low duration avoidance movements; dark blue line, mean ± standard error of mean (SEM) for high duration avoidance movements Dark grey box, baseline period (BL); light grey box, movement period (Move). (G) PETH showing increase in vmPFC-VMS calcium signal at ITI movement initiation for high and low duration ITI movements. Grey line, mean ± standard error of mean (SEM) for low duration ITI movements; black line, mean ± standard error of mean (SEM) for high duration ITI movements Dark grey box, baseline period (BL); light grey box, movement period (Move). (H) Quantification of PETH shows no significant difference in vmPFC-VMS calcium signal between high and low duration movements for both avoidance and ITI movements during the baseline period (−10 to −8s) or the movement period (0 to 1s). (Two-way ANOVA, Task Period x Velocity p = 0.1002, Task Period p < 0.0001, Duration p = 0.3532, Sidak’s Multiple Comparisons Test, High Duration Avoidance Baseline vs Low Duration Avoidance Baseline p = 0.9570, High Duration Avoidance Movement vs Low Duration Avoidance Movement p = 0.3472, High Duration ITI Baseline vs Low Duration ITI Baseline p > 0.9999, High Duration ITI Movement vs Low Duration ITI Movement p = 0.2340, N = 9 mice, High Duration Avoidance Movement n = 127 trials, Low Duration Avoidance Movement n = 110 trials, High Duration ITI Movement n = 704 occurrences, Low Duration ITI Movement n = 480 occurrences). (I) Distribution of movement velocities for avoidance (green) and ITI (black) movements for BLA-VMS data. Green dashed line represents cutoff (10.5 cm/s) for separating avoidance movements of high and low velocity. Black dashed line represents cutoff (2.5cm/s) for separating ITI movements of high and low velocity. (J) PETH showing increase in BLA-VMS calcium signal at avoidance movement initiation for high and low velocity avoidance movements. Light green line, mean ± standard error of mean (SEM) for low velocity avoidance movements; dark green line, mean ± standard error of mean (SEM) for high velocity avoidance movements Dark grey box, baseline period (BL); light grey box, movement period (Move). (K) PETH showing increase in BLA-VMS calcium signal at ITI movement initiation for high and low velocity ITI movements. Grey line, mean ± standard error of mean (SEM) for low velocity ITI movements; black line, mean ± standard error of mean (SEM) for high velocity ITI movements Dark grey box, baseline period (BL); light grey box, movement period (Move). (L) Quantification of PETH shows no significant difference in BLA-VMS calcium signal between high and low duration movements for both avoidance and ITI movements during the baseline period (−10 to - 8s) or the movement period (0 to 1s). (Two-way ANOVA, Task Period x Velocity p = 0.7084, Task Period p = 0.0213, Velocity p = 0.4356, Sidak’s Multiple Comparisons Test, High Velocity Avoidance Baseline vs Low Velocity Avoidance Baseline p > 0.9999, High Velocity Avoidance Movement vs Low Velocity Avoidance Movement p = 0.7110, High Velocity ITI Baseline vs Low Velocity ITI Baseline p = 0.9939, High Velocity ITI Movement vs Low Velocity ITI Movement p = 0.9936, N = 7 mice, High Velocity Avoidance Movement n = 98 trials, Low Velocity Avoidance Movement n = 81 trials, High velocity ITI Movement n = 571 occurrences, Low Velocity ITI Movement n = 419 occurrences). (M) Distribution of movement durations for avoidance (green) and ITI (black) movements for BLA-VMS data. Green dashed line represents cutoff (2.5 s) for separating avoidance movements of high and low duration. Black dashed line represents cutoff (0.5 s) for separating ITI movements of high and low duration. (N) PETH showing increase in BLA-VMS calcium signal at avoidance movement initiation for high and low duration avoidance movements. Light green line, mean ± standard error of mean (SEM) for low duration avoidance movements; dark green line, mean ± standard error of mean (SEM) for high duration avoidance movements Dark grey box, baseline period (BL); light grey box, movement period (Move). (O) PETH showing increase in BLA-VMS calcium signal at ITI movement initiation for high and low duration ITI movements. Grey line, mean ± standard error of mean (SEM) for low duration ITI movements; black line, mean ± standard error of mean (SEM) for high duration ITI movements Dark grey box, baseline period (BL); light grey box, movement period (Move). (P) Quantification of PETH shows no significant difference in BLA-VMS calcium signal between high and low duration movements for both avoidance and ITI movements during the baseline period (−10 to −8s) or the movement period (0 to 1s). (Two-way ANOVA, Task Period x Velocity p = 0.8734, Task Period p = 0.0170, Duration p = 0.4198, Sidak’s Multiple Comparisons Test, High Duration Avoidance Baseline vs Low Duration Avoidance Baseline p = 0.9995, High Duration Avoidance Movement vs Low Duration Avoidance Movement p = 0.8288, High Duration ITI Baseline vs Low Duration ITI Baseline p > 0.9999, High Duration ITI Movement vs Low Duration ITI Movement p = 0.9974, N = 7 mice, High Duration Avoidance Movement n = 95 trials, Low Duration Avoidance Movement n = 87 trials, High Duration ITI Movement n = 585 occurrences, Low Duration ITI Movement n = 413 occurrences). ns = not significant, * p **≤** 0.05.

**Extended Data Figure 6-1.**
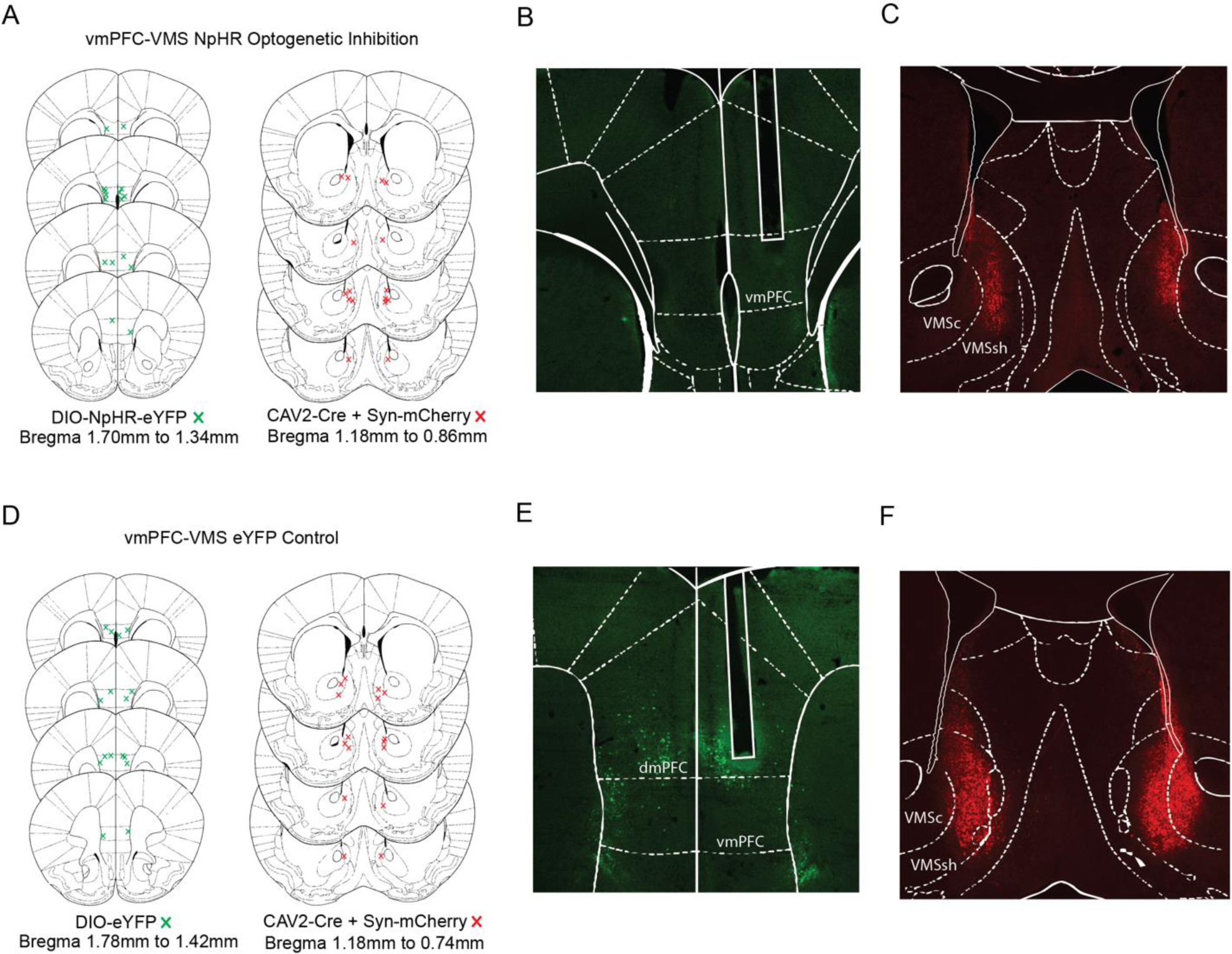
Histology and targeting for vmPFC-VMS optogenetic inhibition surgeries. (A) Verification of bilateral NpHR viral injections in vmPFC (left) and bilateral CAV2-Cre + mCherry viral injections in VMS (right) for the NpHR optogenetic inhibition group (N = 8 mice). (B) Representative histological image of optogenetic implant and NpHR viral expression in vmPFC for the NpHR optogenetic inhibition group. (C) Representative histological image of CAV2-Cre + mCherry viral expression in VMS for the NpHR optogenetic inhibition group. (D) Verification of bilateral eYFP viral injections in vmPFC (left) and bilateral CAV2-Cre + mCherry viral injections in VMS (right) for the eYFP control group (N = 8 mice). (E) Representative histological image of optogenetic implant and eYFP viral expression in vmPFC for the eYFP control group. (F) Representative histological image of CAV2-Cre + mCherry viral expression in VMS for the eYFP control group.

